# Parietal cell-Specific SLC26A9 Deletion induces spontaneous Gastric Carcinogenesis in Mice

**DOI:** 10.1101/2020.09.28.316398

**Authors:** Xuemei Liu, Taolang Li, Dumin Yuan, Brigitte Riederer, Zhiyuan Ma, Jiaxing Zhu, Yunhua Li, Jiaxing An, Guorong Wen, Hai Jin, Chunli Hu, Minglin Zhang, Xiao Yang, Ursula Seidler, Biguang Tuo

## Abstract

Previous study showed that Slc26a9 loss impairs parietal cell function and survival. We investigated whether Slc26a9 loss causes spontaneous gastric carcinogenesis in mice and plays a role in the development and progression in human gastric cancer (GC). Gastric histopathology and potential molecular mechanism were explored in Slc26a9 knockout mice and wild-type littermates as well as Slc26a9fl/fl/Atp4b-Cre and Slc26a9fl/fl mice from 8 days to 18 months by histological and immunohistochemical analyses, quantitative PCR, in situ hybridization, and RNA microarray analysis, respectively. We demonstrated that loss of parietal cell expression of Slc26a9 is the key event to induce spontaneous gastric carcinogenesis in mice, and clarified the sequence of events leading to malignant transformation, including Slc26a9 deficiency in parietal cells resulted in dysregulated differentiation of stem cells in an inflammatory environment, activated Wnt signaling pathway to induce gastric epithelia cell hyperproliferation and apoptosis inhibition, as well as spontaneous epithelial to mesenchymal transition-induced cancer stem cell phenotypes. Downregulation of SLC26A9 correlated with GC patient’s short survival.

**Graphical Abstract:** Loss of parietal cell expression of Slc26a9 is the key event to induce spontaneous gastric carcinogenesis in transgenic mice.

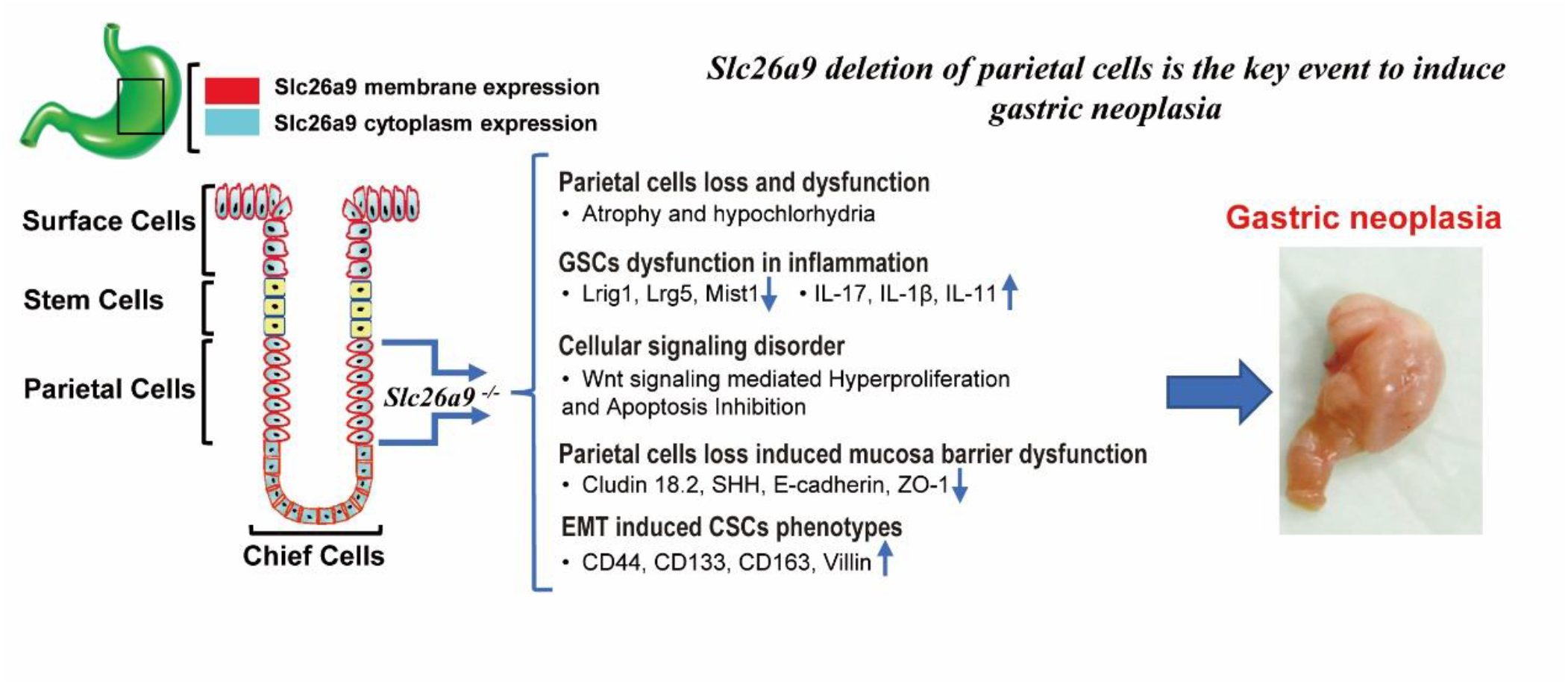

## Introduction

Gastric cancer (GC) is a common malignant tumor and a major health threat in the world. Understanding the critical molecular factors that drive gastric tumor initiation and progression contributes to develop effective strategies for GC prevention and treatment. Gastric carcinogenesis is believed to result from interaction between *Helicobacter pylori (H. pylori)* infection and genetic, epigenetic and environmental factors [1, 2]. It commonly develops through a multistep process of histological progression from atrophic gastritis (AG) through intestinal metaplasia (IM) to GC. In this classic “Correa sequence”, AG with parietal cell loss is a critical initial step necessary for GC development [1, 2], and during the course of multistep carcinogenesis, various genetic and epigenetic alterations accumulate, which facilitates understanding the pathogenesis of GC.

Slc26a9 (solute carrier family 26 member 9) is a member of Slc26a family of multifunctional anion transporters [3], and strongly expressed in gastric surface and glandular epithelium in both mice and humans [4-7]. A previous study showed that deletion of Slc26a9 expression in the murine stomach resulted in progressive parietal cell loss, massive fundic hyperplasia, hypochlorhydria and hypergastrinemia starting a few weeks after birth, suggesting that the Slc26a9 gene is essential for parietal cell function and survival [4]. Parietal cell loss may contribute to the generation of a premalignant environment with mucous cell metaplasia [8]. We therefore wondered whether Slc26a9 gene deficiency may promote gastric carcinogenesis in mice and its mechanisms.

Based on two mouse models, Slc26a9 full knockout mice and parietal cell–specific deletion of Slc26a9 mouse model, we showed that loss of Slc26a9 resulted in spontaneous gastric carcinogenesis in the mice. Slc26a9 deficiency in parietal cells resulted in dysregulated differentiation of stem cells in an inflammatory environment, activated Wnt signaling pathway to induce gastric epithelia cell hyperproliferation and apoptosis inhibition, as well as spontaneous epithelial to mesenchymal transition (EMT) -induced cancer stem cell (CSC) phenotypes. We then studied the expression of SLC26A9 in human GC tissues, and showed that loss of Slc26a9 occurs during the development and progression of human GC and correlates with poor prognosis. Finally, the effect of SLC26A9 on GC cell function was assessed.

## Materials and methods

Please refer to the Supplementary Materials for detailed additional method descriptions.

### Animals

The Slc26a9-deleted mouse strain, whose establishment and characteristics have been described elsewhere[4, 5], was congenic on the S129/svj background. The *Slc26a9*^*fl/fl*^ mice in a C57BL/6J genomic background were obtained from Cyagen Biosciences, China, and *Atp4b-Cre* mice[9] was donated by Prof. Xiao Yang from State Key Laboratory of Proteomics, China. *Slc26a9*^*fl/fl*^ mice were crossed with *Atp4b-Cre* to produce the parietal cells-specific Slc26a9 knockout in *Slc26a9*^*fl/fl*^ */Atp4b-Cre* mice in a mixed genomic background. Mice were bred and genotyped in accordance with the Institutional Animal Care and Use Committee (IACUC) at Hannover Medical School, Germany and at Zunyi Medical University, China, respectively. All mice from 8 days to 18 months of age were age- and sex-matched and used. All experiments involving animals were approved by the Hannover Medical School and Zunyi Medical University committees on investigation involving animals and an independent committee assembled by the local authorities.

### Tissue Microarray and Human Samples

Tissue microarray containing 90 cases of multiple human GC tissues (HStm-Ade180Sur-06), were obtained from Shanghai Outdo Biotech. Human samples including normal gastric epithelium (*H. pylori*-negativity was confirmed by histologic examination for *H. pylori* and C14 urea breath test) and chronic atrophic gastritis (CAG) were obtained in the endoscopy center, 145 GC tissues were collected in the Department of Pathology, Affiliated Hospital of Zunyi Medical University from January 2010 to January 2019. The study was in accordance with the Second Helsinki Declaration and was approved by the Human Subject Committee in Affiliated Hospital of Zunyi Medical University, Zunyi, China. All patients whose biopsies were taken had given written informed consent.

### Statistical Analysis

Data were analyzed by one-or two-way ANOVA using SPSS 19.0 software. The significance of differences was assessed using two-tailed Student’s t-test. The relationship between SLC26A9 expression and the clinical pathologic parameters was examined using Pearson χ2 test. The correlation between SLC26A9 expression and overall survival curves was assessed using Kaplan-Meier plots and compared with the log-rank test. Univariate and multivariate Cox regression analyses were used to evaluate survival data. Significance was set at \**P* < 0.05, \*\**P* <0.01, \*\*\**P* < 0.001; or \*\*\*\**P* < 0.0001; NS, not significantly different.

## Results

### Genetic Deletion of Slc26a9 Leads to Spontaneous Premalignant and Malignant Lesions in Murine Gastric Mucosal Epithelia

Gastric mucosal histopathology was investigated in *slc26a9*^*−/−*^ and wildtype littermates, aged 8 days to 18 months, by using a murine gastric histopathology scoring system published by Rogers [10]. The histopathology scores for heterozygous mice were nearly identical to those of control mice. The gastric mucosae of *slc26a9*^−/−^ were not significantly different from wildtype mice (WM) at 8 days after birth; loss of parietal cells was observed by 1 month of age, oxyntic atrophy (parietal cell and chief cell loss) with elongated and dilated glands was observed by 2 months, and mucous cell metaplasia, including spasmolytic polypeptide-expressing metaplasia (SPEM) and IM, was observed at 6 months. By 14 months after birth, all of the Slc26a9-deficient mice (23/23) exhibited a severe gastric preneoplastic phenotype, including CAG, mucous cell metaplasia, profound cyst formation, and high-grade intraepithelial neoplasia (HGIN) which was considered early GC [10, 11]. Moderately differentiated gastric carcinoma (4/13) and HGIN (9/13) were found in the gastric corpus at 18 months (Figure 1*A-C*), but no submucosal invasion, lymph node or distant metastases or tumor formation in other organs were observed (data not shown).

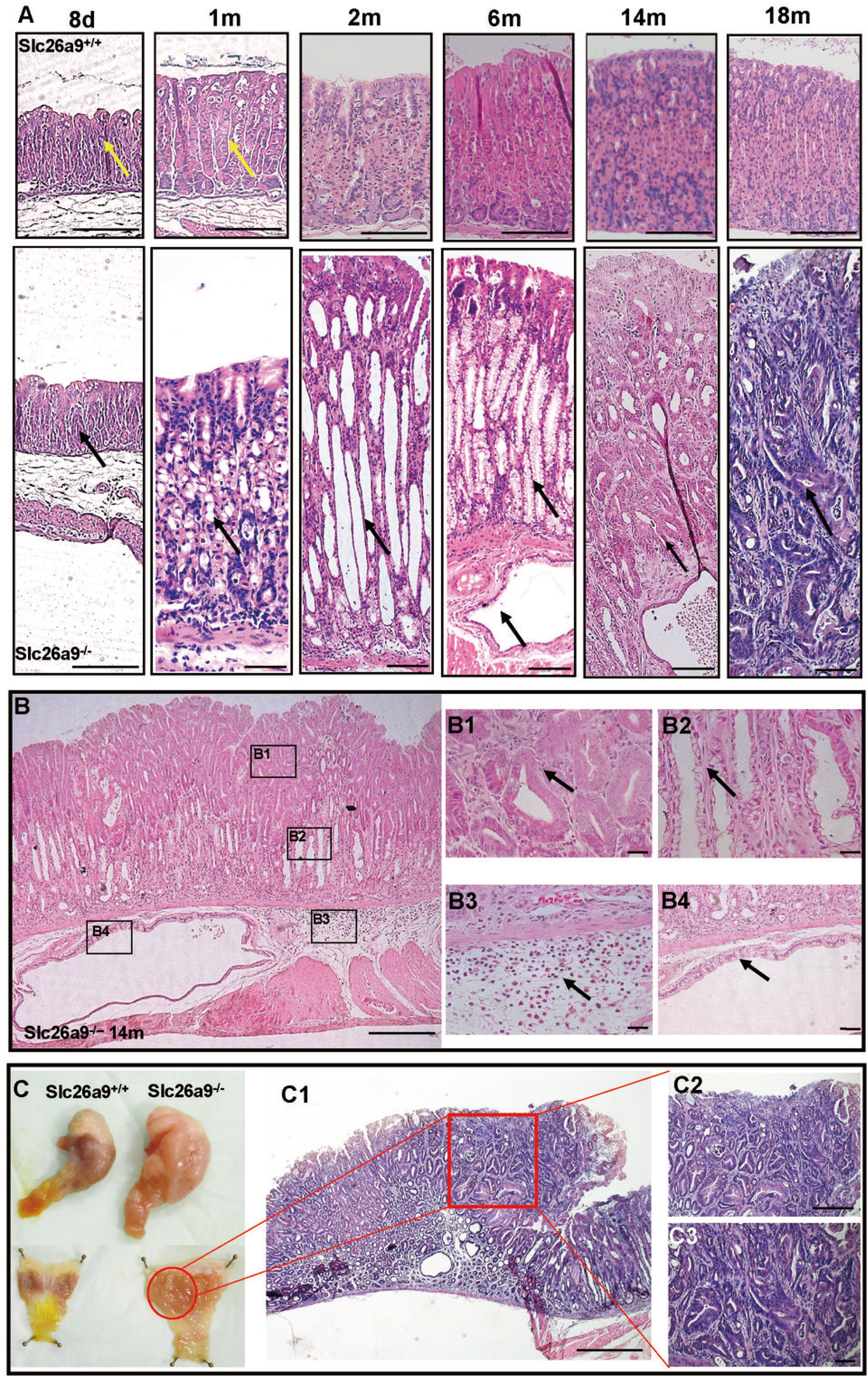

### SLC26A9 Expression is Progressively Downregulated from CAG to GC in Human Gastric Mucosa, and Associated with Poor Prognosis in GC

We next explored the expression of SLC26A9 in human GC. First, qRT-PCR demonstrated that the expression levels of Slc26a9 in human GC tissues were significantly lower than those in normal gastric tissues (Figure 2*A*). Furthermore, IHC analysis showed that SLC26A9 was predominantly localized in the cytoplasm and membrane of surface, parietal, and chief cells in normal gastric epithelia (Figure 2*B*), and it was progressively decreased from CAG to HGIN, and to all types of GC (Figure *2C*), demonstrating that loss of SlC26A9 is associated with the development of GC, which is consistent with our animal experiments.

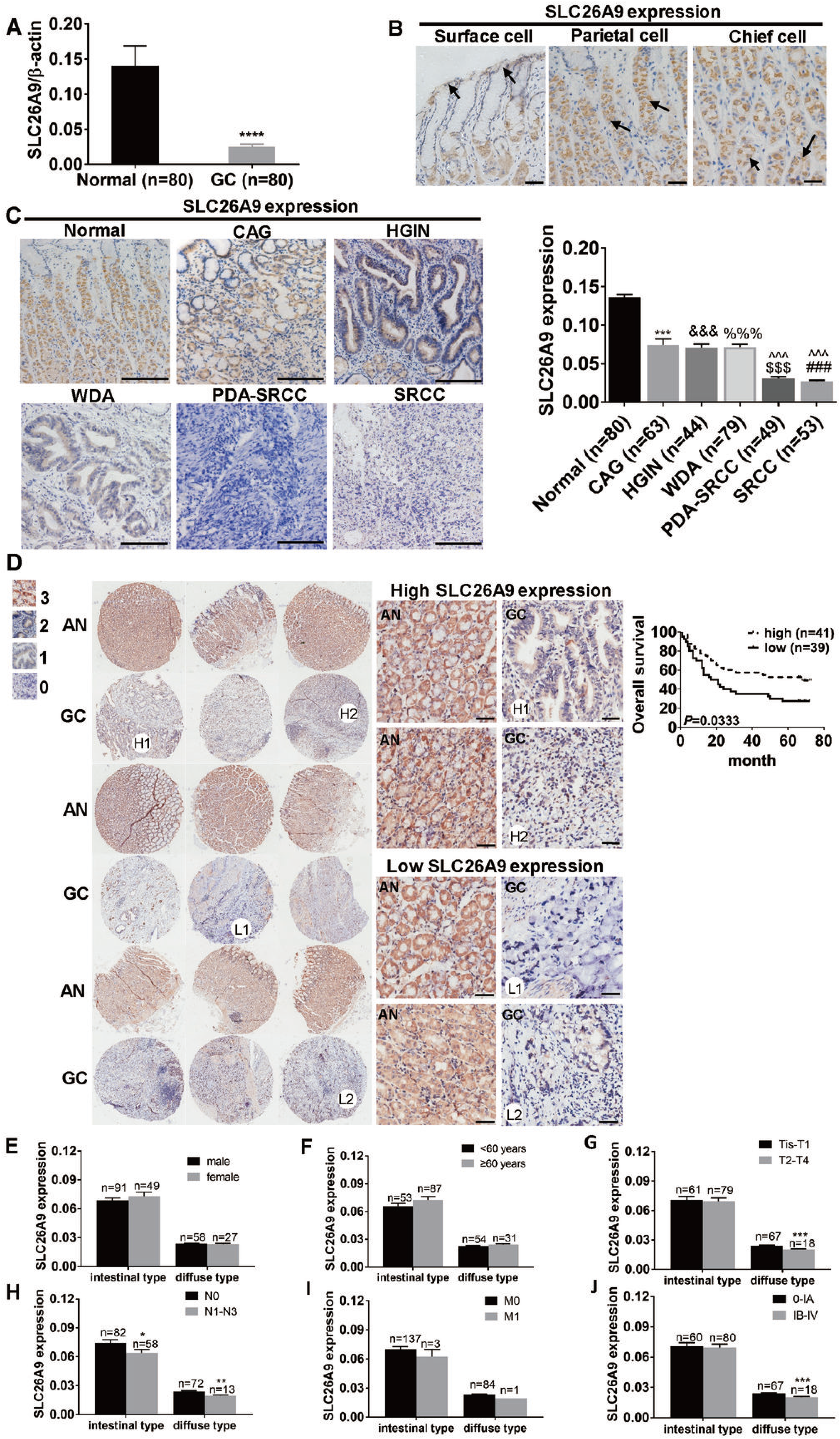

Furthermore, we investigated the relationship between SLC26A9 expression levels in GC and patient’s survival by IHC analysis of a tissue microarray containing 90 GC tissues and 90 adjacent matched normal, of each group 80 tissues were successfully stained and evaluated. After quantifying the SLC26A9 protein expression in the tumor and adjacent normal tissue sections, the patient population was split based on the SLC26A9 IHC staining score (low: score ≤ 1; high: score ≥ 2), revealing the patients in SLC26A9 low expression group had lower overall survival than those in SLC26A9 high expression group (Figure 2*D*).

Finally, we further analyzed SLC26A9 expression in diffuse- and intestinal-type GC tissues of 225 GC patients, including 80 GC samples of tissue microarray and 145 GC tissues from Department of Pathology in our hospital, based on two histological characteristics of GC. The results showed that the expression level of SLC26A9 in GC inversely correlated with its differentiated state (Figure 2*C*). SLC26A9 deficiency was associated with the tumor stage, lymph node stage, metastasis stage, and TNM stage of GC, but not with age or gender in the diffuse type group (Figure 2*E-J*) which consisted of pure signet ring carcinoma and poor differentiated adenocarcinoma with signet ring carcinoma. However, these differences were not observed in the intestinal type GC which included HGIN, well differentiated adenocarcinoma and moderate differentiated adenocarcinoma (Figure 2*E-J*). These results demonstrated that loss of SlC26A9 is associated with a more aggressive phenotype of GC.

### Identification of Molecular Markers for Atrophy, SPEM, IM, Mucosal Barrier Defect, Dysplasia and GC in Slc26a9-deficient Gastric Mucosa

The expression levels of differentiated gastric epithelial cell markers were significantly reduced in the gene panel, and numerous markers associated with both mucous cell metaplasia, including SPEM and IM, mucosal barrier defect and dysplasia, were upregulated in Slc26a9 KM based on transcriptome analysis (Figure 3*A*). Multiple markers specific for CAG, metaplasia, mucosal barrier defect and dysplasia were detected by IHC at 6 months after birth. Compared with WM, Slc26a9 KM exhibited significantly decreased expression levels of gastric mucosal parietal cell markers H^+^/K^+^-ATPase β and Sonic hedgehog (Shh), the chief cell marker mist1 (Figure 3*C*), as well as mucosal barrier defect markers, including foveolar epithelia marker MUC5AC and tight junction marker Claudin 18.2 (Figure 3*F*), but remarkably upregulated levels of specific SPEM markers, including TFF2 and MUC6, from the basement expanding to the upper gastric gland (Figure 3*D*). Moreover, IM was confirmed by overexpression of MUC2, and by finding Alcian blue/Periodic Acid-Schiff-stained cells at the surface and base of gland (Figure 3*E*). These results demonstrated that Slc26a9 KM aged 6 months exhibited robust expression levels of the markers for chronic atrophic gastric, metaplasia, mucosal barrier defect and dysplasia that were also found in human GC [12]. Based on gene microarray analyses, known oncogenes in GC, including ErbB2, Cdkn2a, MAX, TERT, K-Ras, N-Ras, Cyclin E, c-Met and FGFR2, were upregulated at 14 months of age, whereas tumor suppressors, P53, CDH1, and APC, were downregulated (Figure 3*A*). Additionally, the most highly up- and down-regulated genes in Slc26a9 KM were Intelectin 1 (ITLN1) and acidic chitinase (CHIA), respectively (Figure 3*B*). Previous studies demonstrated that the upregulation of ITLN1 is associated with onset of GC in human [13], and the downregulation of CHIA causes gastric atrophy [14]. Moreover, many of the genes from the list of top molecules were highly upregulated, such as CFTR, VNN1 and Cyclin D1 (CCND1), however, CYM, CDKN2A and RunX3 were remarkable downregulated (Figure 3*B*). All of these genes are related to the development of gastric carcinogenesis [15-18].

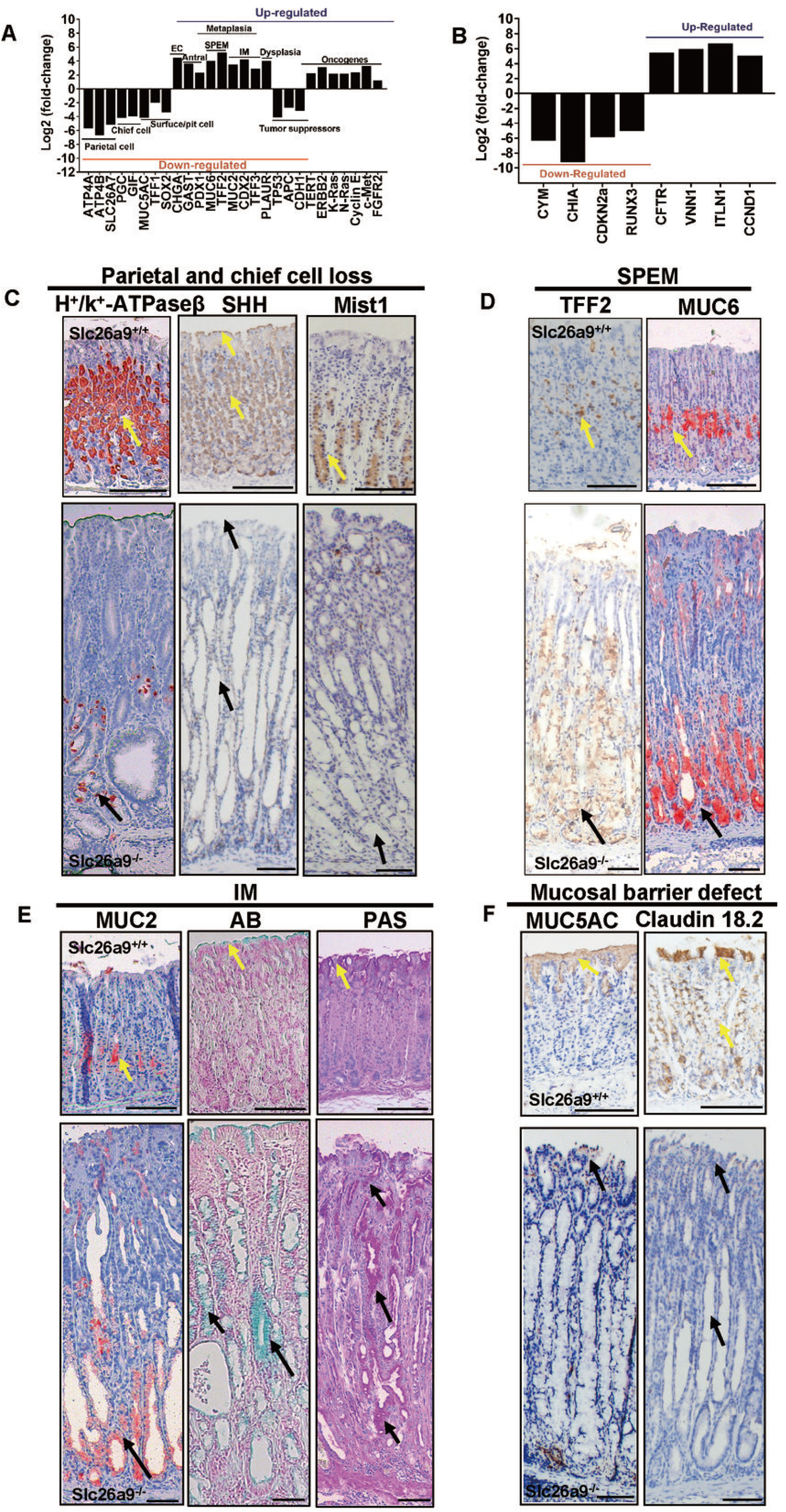

### Slc26a9 Deletion of Parietal Cells is the Key Event to Induce Spontaneous Gastric Carcinogenesis

To further explore whether the initial epithelial dysfunction that leads to parietal cell loss, atrophy, and malignancy is a loss of gastric surface barrier integrity with increased H^+^ back diffusion, or whether the primary event is a defect in parietal cell development, parietal cell-specific Slc26a9 knockout mouse model was established, (Supplementary Figure 1), and gastric mucosal histopathology was investigated in *Slc26a9*^*fl/fl*^*/Atp4b-Cre* and *Slc26a9*^*fl/fl*^ littermates, aged 8 days to 18 months. The gastric mucosae of *Slc26a9*^*fl/fl*^*/Atp4b-Cre* mice were not significantly different from *Slc26a9*^*fl/fl*^ mice at 8 days after birth; loss of parietal cells and oxyntic atrophy with mucous cells metaplasia was observed at 1 month and 6 months, respectively; HGIN was observed in the intromucosa with intact gastric foveolar and surface epithelium at 14 months (Figure 4*A*). Furthermore, compared with *Slc26a9*^*fl/fl*^ mice, *Slc26a9*^*fl/fl*^*/ATP4b-Cre* mice exhibited significantly decreased expression levels of the parietal cell marker H^+^/K^+^-ATPase β in gastric mucosae (Supplementary Figure 2*A*), but with identical expression of foveolar epithelia marker MUC5AC at different time point by IHC analysis (Supplementary Figure 2*B*). Tight junction marker Claudin 18.2 was expressed in the surface cells and parietal cells in the *Slc26a9*^*fl/fl*^ mice, but significantly decreased in the parietal cells and slightly reduced in the surface cells at 14 months, however, no alteration was observed in the gastric surface cells from 8 days to 6 months (Supplementary Figure 2*C*). Nevertheless, poor differentiated carcinoma with mucosal barrier defect was observed in *Slc26a9*^*fl/fl*^*/Atp4b-Cre* mice at 18 months (Figure 4*A*), which was accompanied with loss of MUC5AC and Claudin 18.2 expression when compared with *Slc26a9*^*fl/fl*^ littermates (Figure 4*B*). These data demonstrated that Slc26a9 specific deletion in the parietal cell is the critical step for GC development.

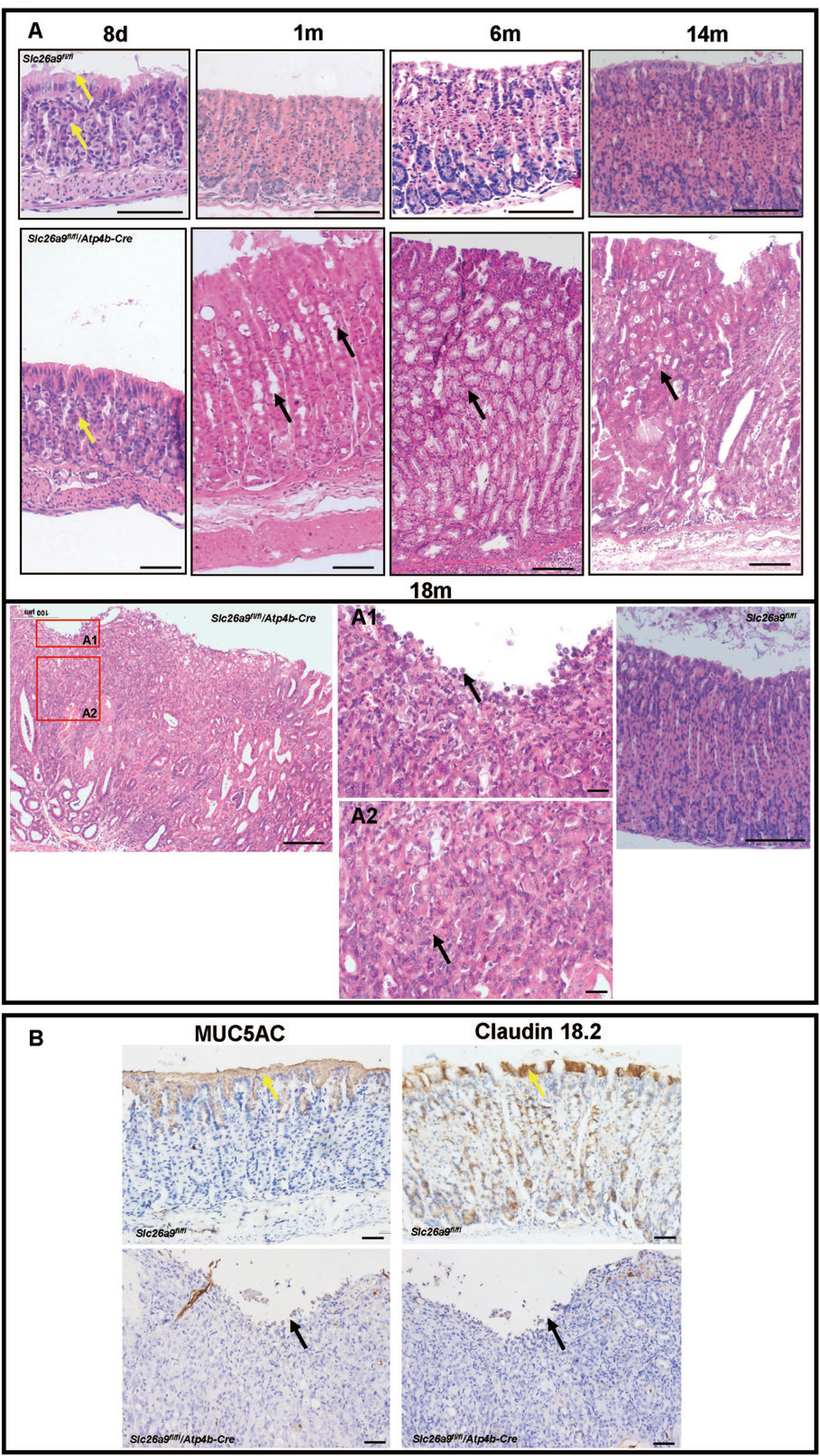

### Slc26a9 Deficiency in the parietal cells Results in Dysregulated Differentiation of Stem and Progenitor cells in an Inflammatory Environment

Loss of Slc26a9 significantly altered not only the markers of gastric epithelial cell differentiation (Figure 3*A*), but also multiple genes encoding ligands, including transforming growth factor-α (TGF-α), amphiregulin (AREG), heparin-binding EGF (HB-EGF), Sonic Hedgehog (SHH), Ptch1, and Notch4 (Figure 5*A*), which are secreted by parietal cells and function as critical regulators of cell differentiation in the gastric mucosa [19], accompanied by inflammatory cytokines IL-17, IL-11 and IL-1β upregulation, indicating spontaneous inflammation (Figure 5*B*). Gastric stem cell (GSC) markers, including Lrig1, Lgr5 and Mist1, were downregulated in *Slc26a9*^*fl/fl*^*/ATP4b-Cre* mice as determined by transcriptome analysis (Figure 5*A*), which was confirmed by RNA in situ hybridization (ISH) and IHC at 1 month, 6 months, and 14 months after birth (Figure 5*C-E*). Previous studies showed that Lrig1 was expressed in both stem cells and parietal cells, implying possible roles for the Lrig1 protein in both cell types [20]. Lgr5, which functions as a stem cell marker of chief cell differentiation, was found to be restricted to a subpopulation of non-proliferative zymogenic chief cells at the base of corpus gland [21, 22]. Mist1 was expressed in both chief cells and stem cells [23]. Our data showed the same expression and localization patterns of Lrig1, Lgr5 and Mist1 in *Slc26a9*^*fl/fl*^ as those reported, but these factors progressively decreased in *Slc26a9*^*fl/fl*^*/ATP4b-Cre* at different time points.

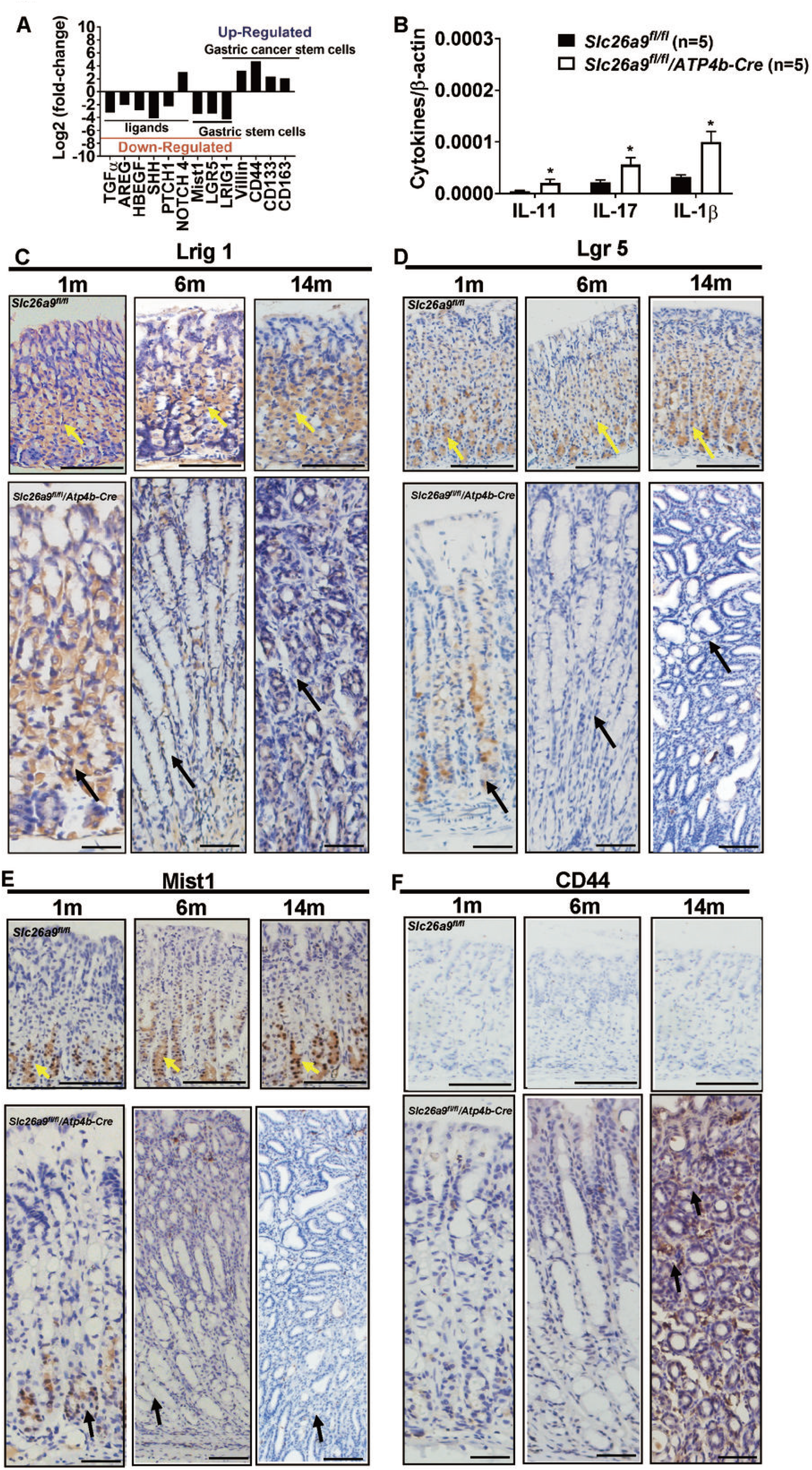

### Slc26a9 Deficiency in Parietal Cells Activated Wnt Signaling pathway to Induce Hyperproliferation and Apoptosis Inhibition

In the gastric mucosa of *Slc26a9*^*fl/fl*^*/ATP4b-Cre* at 14 months of age, the upregulation of 494 genes and downregulation of 642 genes were found by RNA gene microarray analysis. The first 10 signaling pathways and biological processes with *P* values < 0.05 are shown in Figure 6*A* and *B*. These factors were shown to be associated with cell proliferation, differentiation, apoptosis, gastric stem cells, transport activity, digestion, cell and organ development, and homeostasis as well as signal transduction, transcription, and metabolism (Figure 6*A* and *B*, Supplementary Table 3), such as the pathways involving P53, tight junction, SHH, extracellular matrix (ECM) interaction, MAPK, transforming growth factor-β (TGF-β), cell cycle, Wnt, focal adhesion as well as the alteration of digestion- and metabolism-related numerous gene families.

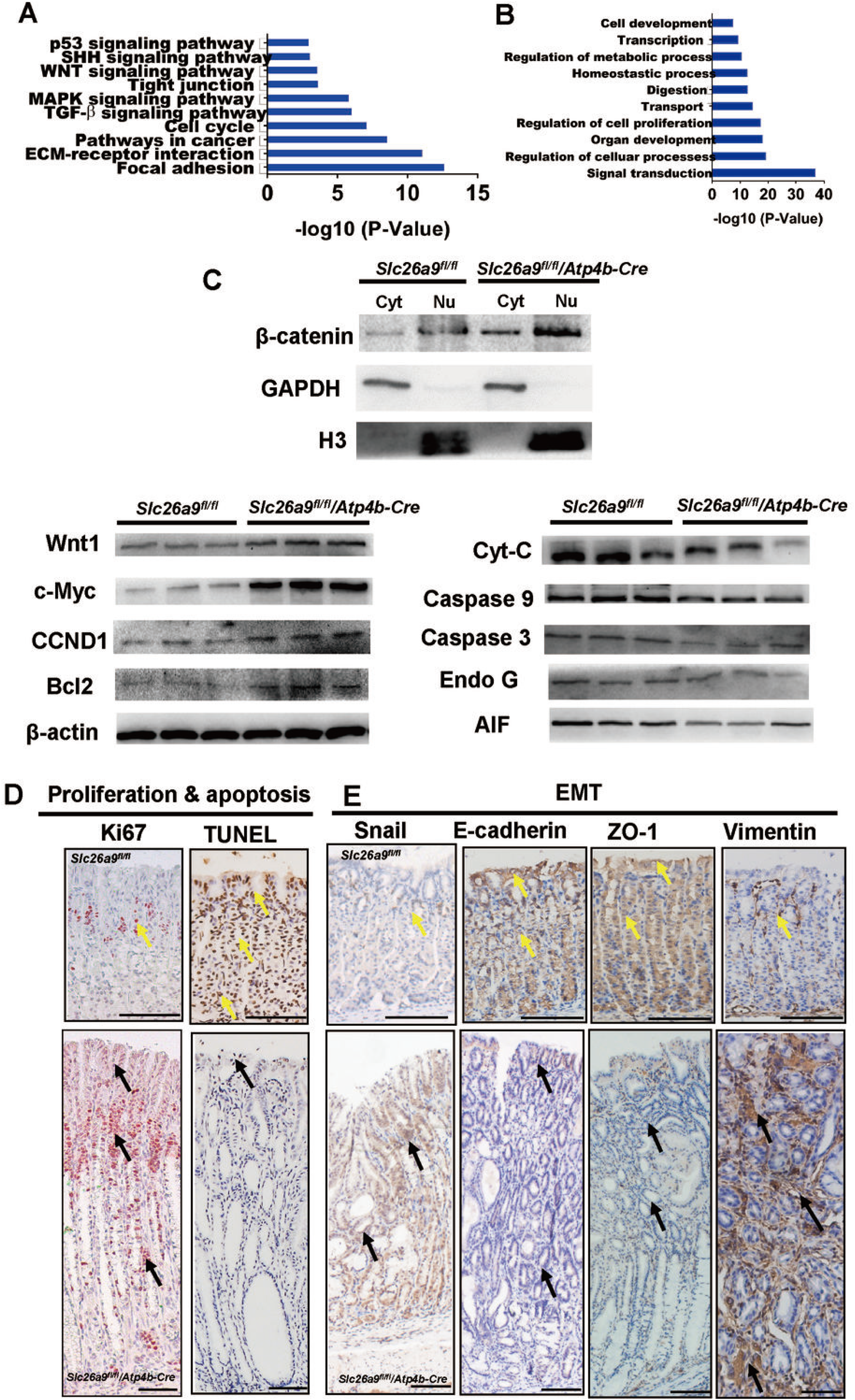

Imbalance of gastric epithelial cell proliferation and apoptosis results in the development of GC, which is highly regulated by Wnt signaling pathway [24]. Generally, translocation of β-catenin to the nucleus reflects the activation of the Wnt pathway and promotes the transcription of downstream targets CCND1, c-Myc and Bcl2 that is associated with hyperproliferation and apoptosis suppression and apoptosis inhibition [24]. Our data showed that expression of Wnt1, β-catenin, and downstream target genes including CCND1, c-Myc and Bcl2 was significantly elevated in the *Slc26a9*^*fl/fl*^*/ATP4b-Cre* mice relative to controls, which resulted in downregulation of cytochrome C (Cyt-C), cleaved Caspase9, cleaved Caspase3, apoptosis-inducing factor (AIF) and endonuclease G (Endo G), indicating that the suppression of caspase dependent and independent apoptosis (Figure 6*C*). We confirmed that in control stomach, Ki67 and terminal deoxynucleotidyl transferase dUTP nick end labeling (TUNEL) assays showed that proliferating cells were localized to the isthmus region and apoptotic cells were displayed in the whole gland in control mouse tissues, however, in *Slc26a9*^*fl/fl*^*/ATP4b-Cre mice*, proliferative cell areas had greatly expanded to the entire length of the gastric glands, but apoptotic cells were significantly less numerous and localized to the surface cells (Figure 6*D*).

### Slc26a9 Deficiency in the parietal cells Results in Spontaneous EMT-Induced Cancer Stem Cell (CSC) Phenotypes

We further explored whether Slc26a9 deficiency caused EMT in the stomach at 14 months after birth due to the loss of epithelial characteristics and cell adhesion. In control mice, two specific epithelial markers, E-cadherin and ZO-1, were predominantly expressed in the membranes of surface, parietal, and chief cells. However, the expression levels of these proteins were significantly decreased in the gastric epithelia of *Slc26a9*^*fl/fl*^*/ATP4b-Cre* mice, with the upregulation of snail and the mesenchymal marker Vimentin in gastric mesenchymal cells (Figure 6*E*), suggesting that the loss of Slc26a9 promotes spontaneous EMT. Moreover, alterations in E-cadherin expression subsequently leads to the accumulation of cytoplasmic β-catenin that then translocates to the nucleus and reflects the activation of the Wnt pathway [24] (Figure *6C* and *E*). During the EMT process, epithelial cells dedifferentiate and acquire mesenchymal as well as EMT-induced CSC phenotypes, which contribute to gastric carcinogenesis [25, 26]. We detected enriched gene expression of surface markers including CD44, CD133, CD163, and Villin in *Slc26a9*^*fl/fl*^*/ATP4b-Cre* mice (Figure 5*A*). The expression and progressive upregulation of CD44 were confirmed by IHC at 1 month, 6 months and 14 months after birth in *Slc26a9*^*fl/fl*^*/ATP4b-Cre* mice, whereas no CD44 expression was detectable in *Slc26a9*^*fl/fl*^ mice (Figure 5*F*).

### Re-expression of SLC26A9 Reduced Gastric Tumor Cell Proliferation and Blocked EMT by Inhibiting Wnt Signaling Pathway

To further confirm Slc26a9 specific deletion in the parietal cells of mice caused imbalance of cell proliferation and apoptosis, as well as spontaneous EMT by mediating Wnt pathway activation. Expression and function of SLC26A9 was investigated in the human GC cell lines. Firstly, SLC26A9 was abundantly expressed in the normal gastric epithelial cell line GES-1, but displayed low expression in the GC cell lines, including MKN45, AGS, SGC-7901, MKN28, and KATOIII, in both mRNA and protein levels. Among these GC cells, the lowest expression was observed in the poorly differentiated adenocarcinoma cell line AGS and the signet ring cell carcinoma cell line KATOIII, and there was the highest expression in the well differentiated adenocarcinoma cell line 7901 (Supplementary Figure 3*A*). These data are consistent with the clinical relevance of SlC26A9 to human GC (Figure *2E-J*), and demonstrates that loss of SlC26A9 is associated with a more aggressive phenotype of GC. Furthermore, we examined the role of SLC26A9 in the biological functions of GC cell *in vitro* using AGS cells transfected with lentivirus carrying the SLC26A9 gene fragment or empty vector (Supplementary Figure 3*B*). Overexpression of SLC26A9 in AGS resulted in not only significant reduction of cell growth and total cell number from day 4 to day 6 after transfection (Supplementary Figure 3*C* and *D*), but also decreased the number of Ki67-positive proliferative cells compared with the control groups (Figure 7*A*). To determine whether the antiproliferative effect of SLC26A9 is due to enhanced cell death in these cells, we examined cell cycle as well as early and later stage apoptosis. Re-expression of SLC26A9 arrested the cell cycle at G2/M checkpoint (Figure 7*B*), followed by increased percentages of early- and late-stage apoptotic cells as determined by annexin V-FITC staining and flow cytometry analysis (Figure 7*C*). TUNEL staining confirmed the late stage apoptosis (Supplementary Figure 3*E*). Moreover, AGS cells with re-expressing SLC26A9 exhibited inhibited migration and invasion capabilities compared with control (Figure 7*D* and *E*). These results suggested that overexpression of SLC26A9 promoted apoptosis via inducing G2/M arrest and suppressed the proliferation, migration and invasion of GC cells. Additionally, compared with control, SLC26A9 overexpression in AGS reduced Vimentin and N-Cadherin expression, but increased E-Cadherin and ZO-1 expression, accompanying with inhibited Wnt1, β-catenin, CCDN1 and snail expressions, indicating that SLC26A9 overexpression blocked hyperproliferation and EMT by inhibiting Wnt signaling pathway (Figure 7*F*).

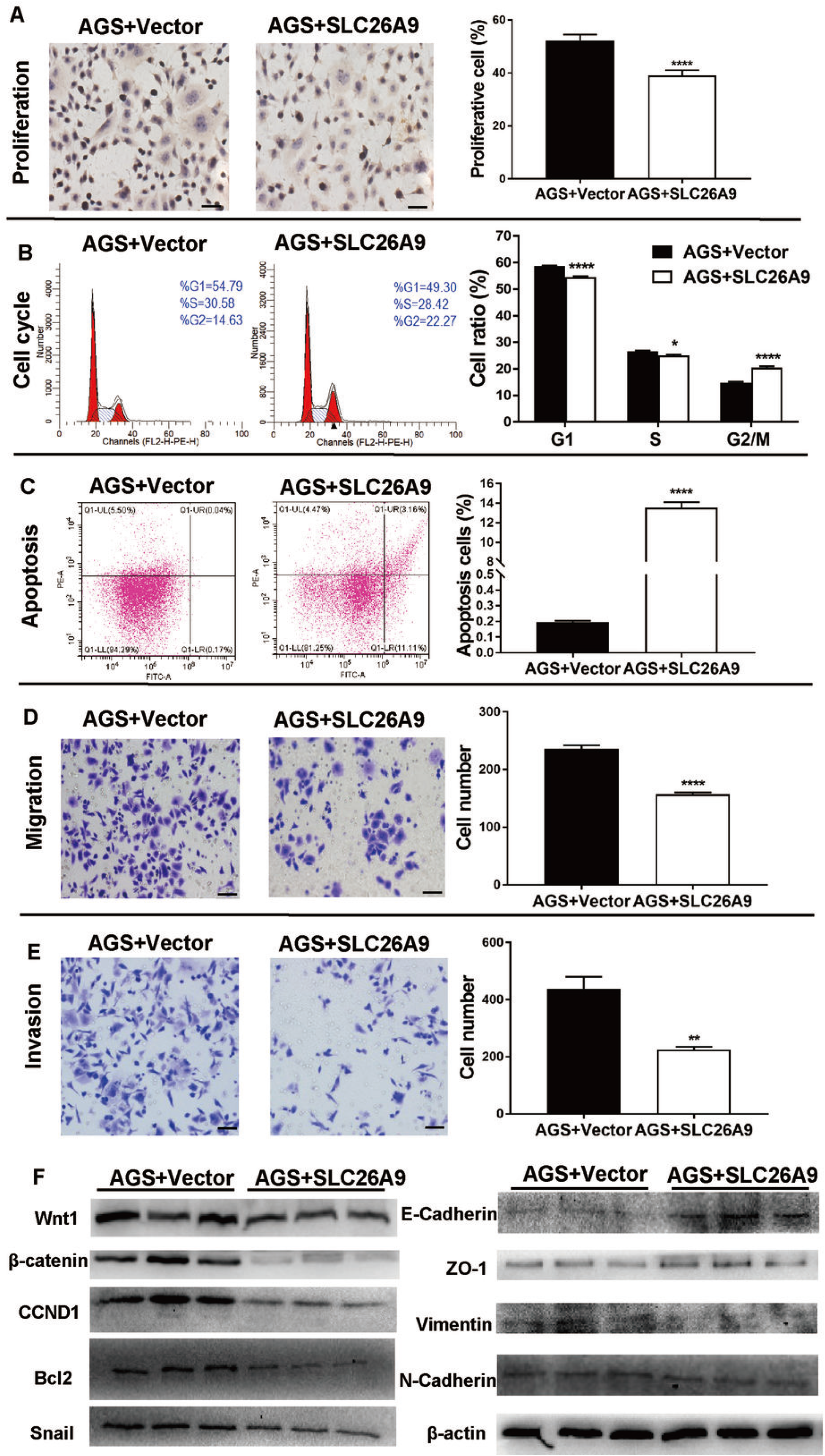

## Discussion

In this study, we demonstrated for the first time that Slc26a9 deficiency causes spontaneous premalignant and malignant lesions in murine gastric epithelia, and clarified the sequence of events leading to malignant transformation.

GC develops through a series of histological changes known as the Correa’s cascade of gastric carcinogenesis. This classic sequence outlines a process of gastric carcinogenesis which is a gradual transition from gastritis to mucosal atrophy, intestinal metaplasia, mucosa barrier defect, dysplasia, and the subsequent development of GC, which is considered to be a classic human model of gastric carcinogenesis [1, 2]. In this study, we showed that Slc26a9 deletion mice displayed histopathologic features from atrophy, SPEM, IM, mucosa barrier defect, dysplasia, finally to GC during 8 days to 18 months after birth. Inflammation, atrophy, IM, and dysplasia were present in Slc26a9 KM aged 14 months. The results from gene microarray and IHC analyses confirmed the molecular phenotypes of atrophy, SPEM, IM dysplasia, and GC in Slc26a9 KM during the development of GC. These results demonstrated that Slc26a9 loss leads to the progress of gastric mucosal epithelial lesions from atrophy, intestinal metaplasia, mucosa barrier defect, dysplasia, finally to GC. Unlike other mouse models of gastric cancer, which rarely spontaneously develop CG through the alteration of a single gene [27], Slc26a9 deletion in mice induced spontaneous development of GC in consistent with the Correa’s cascade of gastric carcinogenesis. The clinical relevance of Slc26a9 to human GC is supported by our findings of progressive SLC26A9 downregulation from CAG to GC in human and SLC26A9 expression level in GC inversely correlated with the differentiated state of GC and patient’s clinical outcome, indicating that SLC26A9 may be involved in GC pathogenesis and progression in humans and it might be a novel poor prognosis marker for GC.

Slc26a9 is an anion transporter that is abundantly expressed in the gastric surface, parietal, and chief cells in both mice and humans, and its deletion results in early loss of gastric acid secretory capacity in mice [4-6]. Based on above results, a crucial question is therefore whether the initial epithelial dysfunction that leads to parietal cell loss, atrophy, and malignancy is a loss of gastric surface barrier integrity with increased H^+^ back diffusion, or whether the primary event is a defect in parietal cell development. Both events have been associated with the development of gastric epithelial metaplasia and increased risk of malignancy in human and murine stomach [8, 28]. We therefore selectively deleted Slc26a9 in cells expressing the β-subunit of the gastric H^+^/K^+^-ATPase, which will be parietal cells precursor [29, 30] and parietal cells. The parietal cell-selective deletion of Slc26a9 resulted in similar histopathologic features and the development of premalignant and malignant phenotype as observed in the stomach of the full knockout. Of note, compared with mucosal barrier defect was observed in Slc26a9 full knockout mice at 6 months after birth (Figure 3*F*), *Slc26a9*^*fl/fl*^*/ATP4b-Cre* mice displayed intact gastric foveolar epithelia and tight junction at same age (Supplementary Figure 2*B* and *C)*. The relative markers for mucosa barrier including MUC5AC and claudin 18.2 were lost when cancerous lesions were detected (Figure 4*B)*. This demonstrates that Slc26a9 expression is highly important for parietal cell function and survival, Slc26a9 specific deficiency in the parietal cells is the critical and initial step to induce spontaneous gastric carcinogenesis.

In this study, loss of parietal cells was observed at 1 month of age in two types of Slc26a9 deletion mouse models, which is considered as the critical and initial step for GC development [31] for the following reasons: First, loss of parietal cell-derived signaling molecules disrupts the proper differentiation of other lineages [8]. We showed that Slc26a9 loss significantly altered not only markers of gastric epithelial cell differentiation, but also multiple genes which encode ligands that are secreted by parietal cells and function as critical growth factors in the gastric mucosa including SHH, Notch4, TGF-α, ARGE and HB-EGF [32]. Second, Slc26a9 deletion resulted in initial parietal and chief cell loss, followed by the emergence of metaplastic lineages including SPEM and IM, which are both known to be associated with an increased susceptibility to neoplastic transformation [31]. Third, impaired acid secretion promotes excessive bacteria growth that triggers the upregulation of related inflammatory factors and leads to intragastric infection and CAG, eventually progressing to GC [33]. Slc26a9 deletion caused hypochlorhydria [4] with an increased expression of the proinflammatory cytokines IL-17 [34], IL-11 [35], and IL-1β [36], which are well known to strongly promote carcinogenesis.

Gastric carcinogenesis is not only a multistep process involving numerous genetic and epigenetic changes in oncogenes, tumor-suppressor genes, cell-cycle regulators, cell adhesion molecules, and DNA repair genes [37], but also resulting from dysregulated differentiation of stem and progenitor cells in an inflammatory environment [2]. We found that Slc26a9 deletion not only disrupted gastric epithelial differentiation and oncogene and tumor-suppressor gene expression associated with GC (Figure 3*A*), but also induced progressive downregulation of GSCs markers, Lrig1, Lgr5, and Mist1, at different time point, accompanying with chronic inflammation, suggesting the disruption of normal gastric differentiation into parietal cells and chief cells, respectively. These data demonstrated the Slc26a9 leads to dysregulated differentiation of stem and progenitor cells in the inflammatory environment to cause gastric carcinogenesis [2]. Furthermore, Slc26a9 deletion leaded to the alteration of multiple signaling pathways that are related to the regulation of cell proliferation, apoptosis, and differentiation, as well as barrier integrity. Emerging evidence suggests that activation of Wnt signaling pathway can be linked to GC development and progression [12, 38]. Activation of the Wnt pathway is known to result in β-catenin nuclear accumulation [24]. Cytoplasmic accumulation and nuclear translocation of β-catenin was observed with increased canonical Wnt protein expression in the *Slc26a9*^*fl/fl*^*/ATP4b-Cre* mice. Accumulated β-catenin in the cytoplasm eventually translocates to the nucleus, which subsequently regulates target proteins including CCND1 and cMyc, which plays an important role in proliferation [24]. Consistent with nuclear expression of β-catenin in the *Slc26a9*^*fl/fl*^*/ATP4b-Cre* mice, increased CCND1, cMyc and ki67 incorporation was reflective of increased cellular proliferation. However, increased Bcl2 with decreased Cyt-C, cleaved Caspase9, cleaved Caspase3, AIF and Endo G, as well as TUNEL incorporation was reflective of suppression of caspase dependent and independent apoptosis [39]. Furthermore, it is known that E-cadherin expression negatively controls the transcriptional activity of β-catenin [24]. Increased Snail expression was accompanied by a significant decrease in E-cadherin expression in the gastric mucosa [40]. Our data showed that dysregulation of surface epithelia and tight junctional polarity protein expression, including MUC5AC, ZO-1, E-cadherin and Claudin 18.2, which was accompanied by upregulation of Snail and the mesenchymal marker Vimentin in gastric mesenchymal cells at 14 months, indicating spontaneous EMT. EMT promotes the generation of cancer stem cells [26] including CD44, CD166, and CD136, especially CD44 overexpression over time. Collectively, loss of Slc26a9 triggers a number of molecular events, including translocation of β-catenin, activation of the Wnt pathway, increased Snail and loss of E-cadherin expression which resulted in gastric epithelia cell proliferation and apoptosis imbalance, as well as EMT-induced tumorigenicity.

The elucidation of the crystal structure of mammalian Slc26a9 revealed that the protein acts as a Cl^−^ uniporter [41], and it is reported to mediate Cl^−^ efflux in parietal [4] and airways cells [42]. Overexpression of SLC26A9 in AGS cells promoted apoptosis via inducing G2/M arrest and suppressed the proliferation, migration and invasion of the GC cells. How can we imagine that a Cl^−^ transporter plays a role in promoting survival in one cell type, while enhancing apoptotic cell death in another? Indeed, anion channels have important but highly diverse functions in cell growth and cell death. By inducing cell shrinkage, they can initiate apoptotic cell death. Identical channels may support regulated death in some cell types, but may cause cell proliferation, cancer development, and metastasis in others [43]. Increased proliferation was also observed in the duodenal crypts of Slc26a9-deficient mice [5]. Furthermore, it is known that EMT induces progression by activation of Wnt signaling [44]. Our study indicated that Slc26a9 suppressed GC cell migration and invasion through attenuating canonical Wnt signaling pathway and the EMT process, which not only further support the findings of animal experiments (Figure 6*C-E*), but also may provide novel therapeutic strategies for GC progression.

## Conclusions

In summary, this study first time demonstrates that Slc26a9 might function as a novel tumor suppressor gene, possibly by maintaining parietal cell viability, normal differentiation of gastric stem cell and Wnt signaling transduction, supporting mucosal homeostasis and acid/base equilibrium in the stomach. Conversely, loss of Slc26a9 in the parietal cells is the key and initial step to induce gastric carcinogenesis in mice and results in the development and progression of human GC and correlates with poor prognosis. These findings provide two novel models of GC in consistent with the Correa’s cascade of gastric carcinogenesis and new insights into the molecular pathogenesis of GC, and SLC26a9 might be important prognostic marker for GC, which contributes to develop interventional strategies for the prevention and treatment of this disease.

## Abbreviations used in this paper

AG: atrophic gastritis
AIF: apoptosis-inducing factor
AREG: amphiregulin
CAG: chronic atrophic gastritis
CCND1: Cyclin D1
CHIA: acidic chitinase
Cyt-C: cytochrome C
EGFR: epidermal growth factor receptor
EMT: epithelial-mesenchymal transition
Endo G: endonuclease G
GC: gastric cancer
GCS: gastric cancer stem cell
GSC: gastric stem cell
HGIN: high-grade intraepithelial neoplasia
HB-EGF: heparin-binding EGF
IM: intestine metaplasia
ISH: in situ hybridization
ITLN1: intelectin 1
KM: knockout mice
SPEM: spasmolytic polypeptide-expressing metaplasia
TGF-α: transforming growth factor-α
TUNEL: transferase dUTP Nick End Labelling
WM: wild type mice

## Authors’ contributions

X.L., T.L., U.S., and B.T. conceived the study concept and design and wrote the manuscript. D.Y., Z.M., C.H., J.Z., G.W., H.J., J.A., X.C., B.R., and B.R. acquired the data. J.Z. assigned the histopathology scores. B.R., B.R., and D.Y. oversaw the mouse studies. All authors read and approved the final manuscript, and X.L., B.T., and U.S. edited the manuscript.

## Acknowledgments

We sincerely appreciate the help of the animal caretakers of the institute of animal research from Zunyi Medical University, China and Hannover Medical School, Germany in breeding and scoring the two types of Slc26a9-deficient mouse strain.

## Ethics approval and consent to participate

Human samples were collected in the Endoscopy Center and Department of Pathology, Affiliated Hospital of Zunyi Medical University. The study was in accordance with the Second Helsinki Declaration and approved by the Human Subject Committee in Affiliated Hospital of Zunyi Medical University, Zunyi, China. All patients whose biopsies were taken had given written informed consent.

